# Re-analysis of SARS-CoV-2 infected host cell proteomics time-course data by impact pathway analysis and network analysis. A potential link with inflammatory response

**DOI:** 10.1101/2020.03.26.009605

**Authors:** Ignacio Ortea, Jens-Ole Bock

## Abstract

The disease known as coronavirus disease 19 (COVID-19), potentially caused by an outbreak of the severe acute respiratory syndrome-coronavirus 2 (SARS-CoV-2) in Wuhan, China, has hit the world hard, and has led to an unprecedent health and economic crisis. In order to develop treatment options able to stop or ameliorate SARS-CoV-2 effects, we need to understand the biology of the virus inside cells, but this kind of studies are still scarce. A recent study investigated translatome and proteome host cell changes induced in vitro by SARS-CoV-2. In the present study, we use the publicly available proteomics data from this study to re-analyze the mechanisms altered by the virus infection by impact pathways analysis and network analysis. Proteins linked to inflammatory response, but also proteins related to chromosome segregation during mitosis, were found to be regulated. The up-regulation of the inflammatory-related proteins observed could be linked to the propagation of inflammatory reaction and lung injury that is observed in advanced stages of COVID-19 patients.

## 1. Introduction

The disease known as coronavirus disease 19 (COVID-19), potentially caused by an outbreak of the severe acute respiratory syndrome-coronavirus 2 (SARS-CoV-2), has hit the world hard^1^. Initially reported in December 2019 in the Chinese city of Wuhan, and potentially linked to a zoonosis related to a wild animal market, COVID-19 has spread globally very rapidly, and World Health Organization (WHO) declared it a pandemic on March 11^th^ 2020. As of March 25^th^ 2020, there are 416,686 confirmed cases and 18,589 confirmed deaths, with 197 countries affected (WHO, www.who.int, data accessed on March 23^rd^ 2020), becoming the biggest health emergency of the 21st century. Because there is currently neither effective treatment nor vaccine, the main measure taken by nations has been social distancing first, and then partial or total preventive lockdown afterwards, deriving in the biggest global economic crisis also of the 21st century.

The typical clinical manifestation of COVID-19 consists of an acute respiratory distress syndrome with fever, dry cough and difficulty breathing, and some patients, especially those with specific comorbidities, can rapidly worsen and die^2,3^, although it is estimated a high proportion of undocumented cases from asymptomatic carriers and mild symptom patients not being tested^4,5^. The crude mortality ratio has been estimated by WHO between 3-4%^6^, although, as a consequence of the high rate of covert cases not being tested, there should be a strong bias on the true mortality rate depending on the number of diagnostic tests performed by each country. In any case, COVID-19 presents alarming levels of spread and severity.

In order to develop treatment options able to stop or ameliorate SARS-CoV-2 effects, we need to understand the biology of the virus inside cells, and therefore there is an urgent need for decipher the host cell molecular mechanisms that are triggered by the virus infection. For instance, cellular factors used by SARS-CoV-2 for the first step of infection, entry into cells, have been recently studied, demonstrating that it employs the angiotensin-converting enzyme 2 (ACE2) host cell receptor, together with the serine protease TMPRSS2, and subsequently a TMPRSS2 inhibitor has been proposed as a treatment option^7^. On the other hand, it has also been reported that ACE2 expression protects from lung injury and is downregulated by SARS-CoV^8,9^, which might promote lung injury, therefore worsening the prognosis of the disease, but it has not been demonstrated yet whether SARS-CoV-2 also interferes with ACE2 expression^7^.

However, the knowledge of what is going on inside the cell after the entry of the virus is still scarce. Host cell proteomics studies, measuring protein abundance changes caused by the virus and obtaining a global vision of these changes by pathway and network analysis, can shed some light on the mechanisms that are used and/or altered by the virus and therefore are targets for drugs to be developed or trialed. To the best of our knowledge, the first available study describing translatome and proteome host cell changes induced by SARS-CoV-2 is the one by Bojkova et al.^10^, where they use Cytoscape and ReactomeFI to propose overrepresented pathways that could be targeted by potential treatment compounds. In this study, we use the publicly available proteomics data from this study to re-analyze the mechanisms altered by the virus infection by impact pathways analysis^11^ and network analysis.

## 2. Materials and Methods

### 2.1. Publicly available proteomics data from cell samples infected with SARS-CoV-2 virus

Proteome measurements from Bojkova et al. (2020)^10^ were downloaded and used for subsequent analysis. Data consisted on the quantification of 6381 proteins in human Caco-2 cell secretomes at four time points after infection with SARS-CoV-2 virus. According to the authors, a TMT-labelling bottom-up quantitative proteomics approach was used to obtain the data, with high pH reverse phase peptide fractionation and mass spectrometry measurement of the peptides using a Thermo QExactive and a nano-liquid chromatography configuration.

### 2.2. Analysis by impact pathway analysis and network analysis

iPathwayGuide (Advaita Corporation, Plymouth, MI, USA) v1910, within the PIPPR pathways analysis framework (COBO Technologies Aps, Maaloev, Denmark), was used for analyzing the significantly impacted pathways and for GO analysis. All quantified proteins were included in the analysis, and the threshold for considering a protein as differentially expressed (DE) was fold-change (log2) higher than 0.5 and p-value below 0.05. Data was analyzed in the context of pathways obtained from the Kyoto Encyclopedia of Genes and Genomes (KEGG) database (Release 90.0+/05-29, May 2019). iPathwayGuide was also used for network analysis, using String v11.0 Jan 2019 and BioGRID v3.5.171 Mar 2019 as data sources. The interactions included were activation, binding, catalysis, expression, and inhibition. Confidence score for protein-protein interaction was set at 900 (high).

### 2.3. Statistical analysis

For impact pathway analysis, iPathwayGuide software calculated a p-value using a hypergeometric distribution. P-values were adjusted using false discovery rate (FDR).

## 3. Results

The affected pathways were analyzed using iPathwayGuide software. Significantly impacted pathways according to this analysis are shown in Figure 1. After FDR correction, six pathways were found to be significantly impacted after 24 h infection, only two at 6 h, and none at 2 h and 10 h (Figure 1a). Expression changes over post-infection time points for selected proteins are shown in Figure 1b.

**Figure 1.**
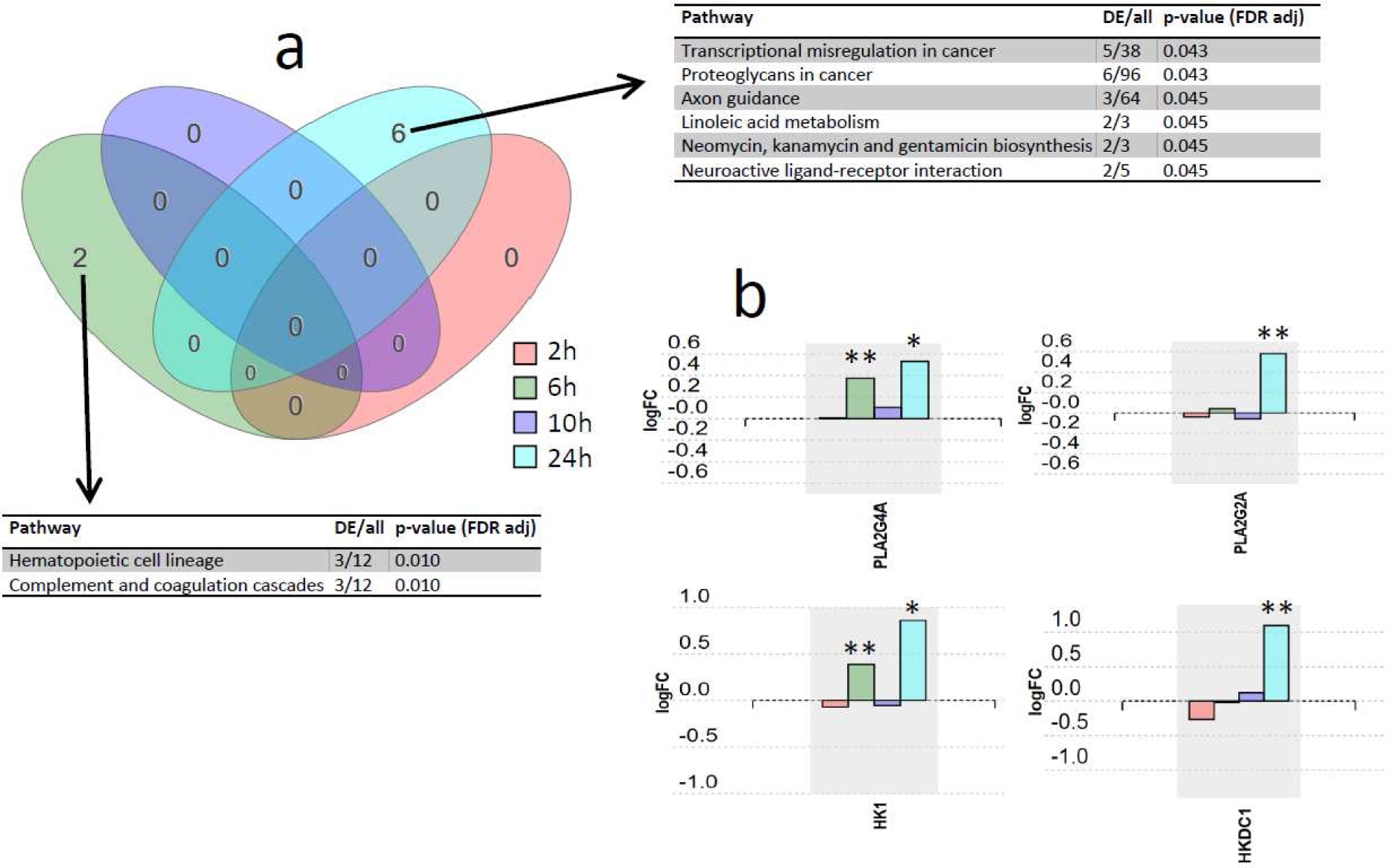
Pathway analysis results. (a) Venn diagram representing the intersections of pathways sets associated to the four post-infection time points. Pathways were considered significant according to a p-value calculated by iPathway Guide software using a hypergeometric distribution and adjusted using false discovery rate. DE, differentially expressed proteins. (b) Expression changes over post-infection time points for proteins PLA2G4A, PLA2G2A, HK1, and HKDC1. * p-value < 0.05, ** p-value < 0.001.

The differentially expressed proteins at the time point presenting the deepest changes (24h after infection), were subjected to network analysis using iPathwayGuide. The interactions included were activation, binding, catalysis, expression, and inhibition. Confidence score for protein-protein interaction was set at 900 (high). The resulting network is shown in Figure 2a. One of the subnetworks with the highest number of interactions, comprised of six proteins, is shown in Figure 2b, together with the expression change profile over post-infection time for these six proteins.

**Figure 2.**
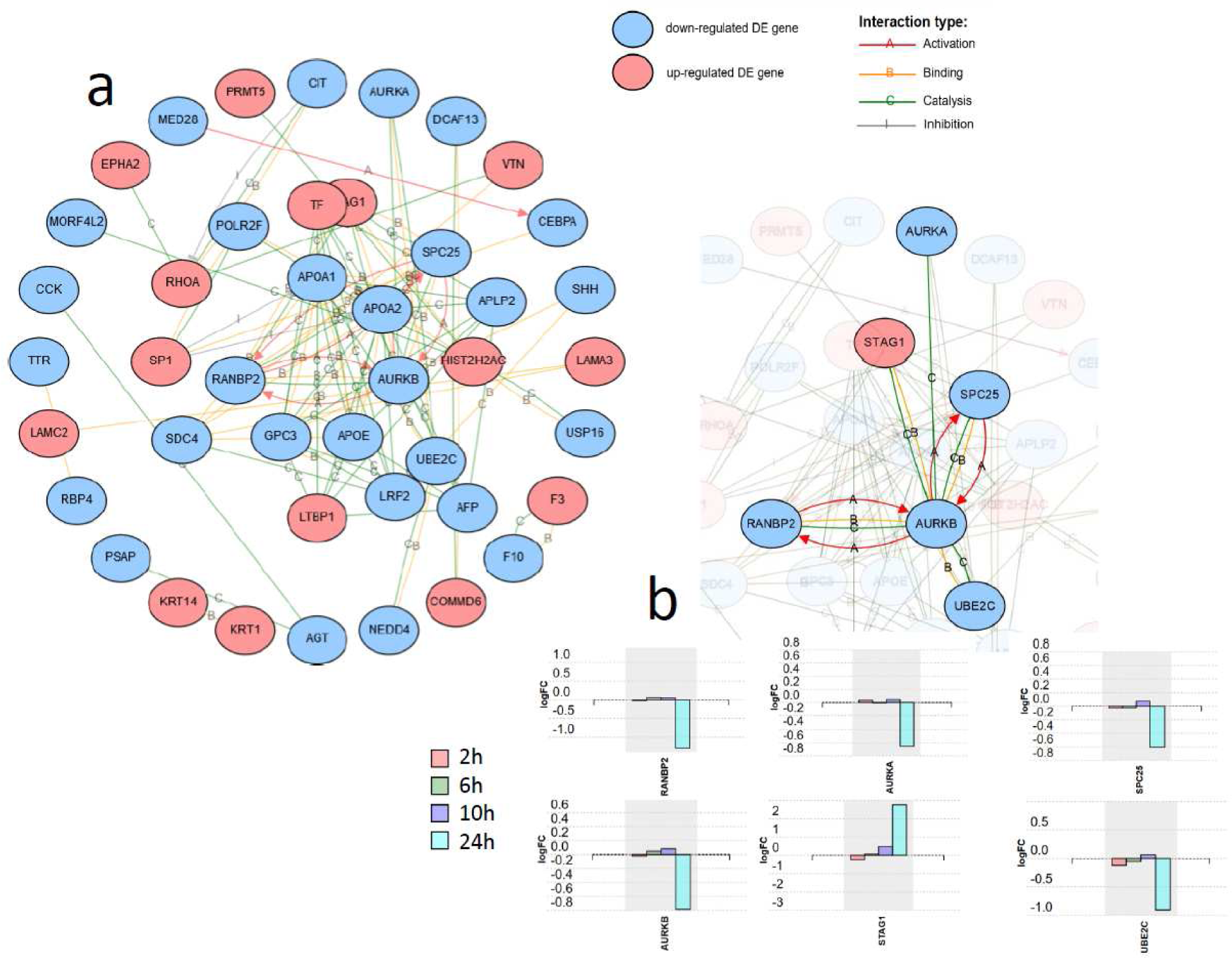
Network analysis including the 125 differentially expressed proteins at 24 h after SARS-CoV-2 in Caco-2 cells. Activation, binding, catalysis, and inhibition regulatory interactions are included. (a) Network with the isolated nodes hidden. (b) Six-protein subnetwork with the interactions for RANBP2, showing the expression changes for each time point for the six proteins.

## 4. Discussion

The significantly affected pathways were analyzed using iPathwayGuide software, which implements an ‘impact analysis’ approach, taking into consideration not only the over-representation of differentially expressed genes in a given pathway (i.e. enrichment analysis), but also topological information such as the direction and type of all signals in a pathway, and the position, role, and type of each protein^11^. Although six pathways were found to be significantly impacted at 24 h, and two at 6 h, the number of DE proteins in these pathways were low (ranging 2 to 6 proteins). For instance, the pathway transcriptional misregulation in cancer, with 5 DE proteins out of the 38 included in the pathway; the proteoglycans in cancer pathway, with 6 DE proteins out of 96; or the axon guidance pathway, with 3 DE proteins out of a total of 64 proteins. Thus, we consider the experimental evidence for having an actual effect of the virus over these mechanisms is relatively low. However, still having a low number of DE proteins, the ratio of DE proteins to total proteins included in three other significant pathways was higher, and for that they deserve a closer look. They are linoleic acid metabolism pathway; neomycin, kanamycin and gentamicin biosynthesis pathway; and neuroactive ligand-receptor interaction pathway. Linoleic acid metabolism pathway is linked to arachidonic acid metabolism and eicosanoids pathway, and therefore it could play a role in the inflammatory response observed in stages II and III of COVID-19 patients^12^. Actually, the two proteins found DE in this pathway at 24 h post-infection, PLA2G4A (cytosolic phospholipase A2) and PLA2G2A (phospholipase A2, membrane associated), are key components of the phospholipase A2 group, which have been previously suggested to participate in a key mechanism in the propagation of inflammatory reaction^13^, and it has been demonstrated its contribution to inflammation and eicosanoid profile in arthritis^14^ and in cardiovascular diseases^15^. When checking the trend over the whole time course, both proteins share the same profile with a clear increase at 24 h after virus infection (Figure 1b). It will then be interesting to explore the potential of these two proteins as COVID-19 early systemic diagnosis biomarkers.

Two proteins from the neomycin, kanamycin and gentamicin biosynthesis pathway resulted up-regulated at 24 h after virus infection (Figure 1b): HK1 (hexokinase 1) and HKDC1 (hexokinase domain containing 1), which are proteins related to glucose use and homeostasis^16,17^. Interestingly, HK has been previously associated with inflammatory response in autoimmune disorders, and, deoxy-D-glucose (2-DG), an inhibitor of HK, has been proposed to ameliorate autoimmune inflammation^18^. Recently, 2-DG has been shown to inhibit SARS-CoV-2 replication in Caco-2 cells^10^ and to inhibit rhinovirus infection and inflammation in a murine model^19^. For all this, hexokinase link to SARS-CoV-2 infection and related inflammation response deserves further study.

In Figure 2a, the network formed by the DE proteins, excluding isolated nodes, is shown. One of the subnetworks with a higher number of connections is the one formed by RANBP2 (E3 SUMO-protein ligase RanBP2) (Figure 2b). RANBP2 forms a complex at the nuclear pore with TRIM5α, a cytoplasmic restriction factor that blocks post-entry retroviral infection and is regulated by SUMO. It has been demonstrated that loss of RANBP2 blocked SUMOylation of TRIM5α, suppressing its anti-retroviral activity^20^. Here RANBP2 presented a statistically significant fold-change (log) of −1.295 at 24 h post-infection, therefore the role of RAMBP2-TRIM5α in coronavirus infection deserves further consideration. In the same subnetwork as RANBP2, some proteins, which are interestingly related to cell cycle progression, also deserve further research (Figure 2b): AURKA, AURKB, SPC25 and STAG1 participate in the regulation of chromosome segregation during mitosis^21-24^. Interestingly, they were all found down-regulated at 24 h post-infection, except STAG1, which was strongly up-regulated. In this subnetwork, closely related to AURKB, and also down-regulated, is UBE2C (Ubiquitin-conjugating enzyme E2 C), which is an essential factor of the anaphase promoting complex/cyclosome (APC/C), a cell cycle-regulated ubiquitin ligase that controls progression through mitosis^25^.

In parallel to re-analyzing the data with alternative tools, when taking a close look at the data we discovered a down-regulation trend over time of ACE2 (Figure 3) that had not been highlighted by the original authors^10^. Actually, at 24 h time ACE2 quantitation presented a fold-change (log) of −0.168 and a p-value of 0.01. Coronavirus entry into target cells depends on binding of its spike (S) proteins to a cellular receptor, which facilitates viral attachment to the surface of target cells. ACE2 was reported as the entry receptor for SARS-CoV^26^, another coronavirus closely related to SARS-CoV-2, playing a key role in SARS-CoV transmissibility^27^, and recently also for by SARS-CoV-2^7^. ACE2 is also a peptidase in the renin-angiotensin system, converting antiotensin I to angiotensin (1-9) and angiotensin II to angiotensin (1-7), a vasodilator. The protective role in lung injury is related to this cleavage of angiotensin II. Regarding ACE2 and SARS-CoV, it was also reported that ACE2 expression protects from lung injury and is downregulated by SARS-CoV^8,9^, which might promote lung injury, therefore worsening the prognosis of the disease. Here we highlight that SARS-CoV2 seems to be also interfering with ACE2 expression, which could be related to a higher level of lung injury as it was demonstrated for SARS-Cov. When inspecting the quantitative data for other proteins in the renin-angiotensin system, two other proteins were found to be down-regulated 24 h post-infection cathepsin A (CTSA) and angiotensinogen (AGT) (Figure 3). We hypothesize that this dysregulation of some of the key components of the renin-angiotensin system could be related to the lung injury and worsening observed in COVID-19.

**Figure 3.**
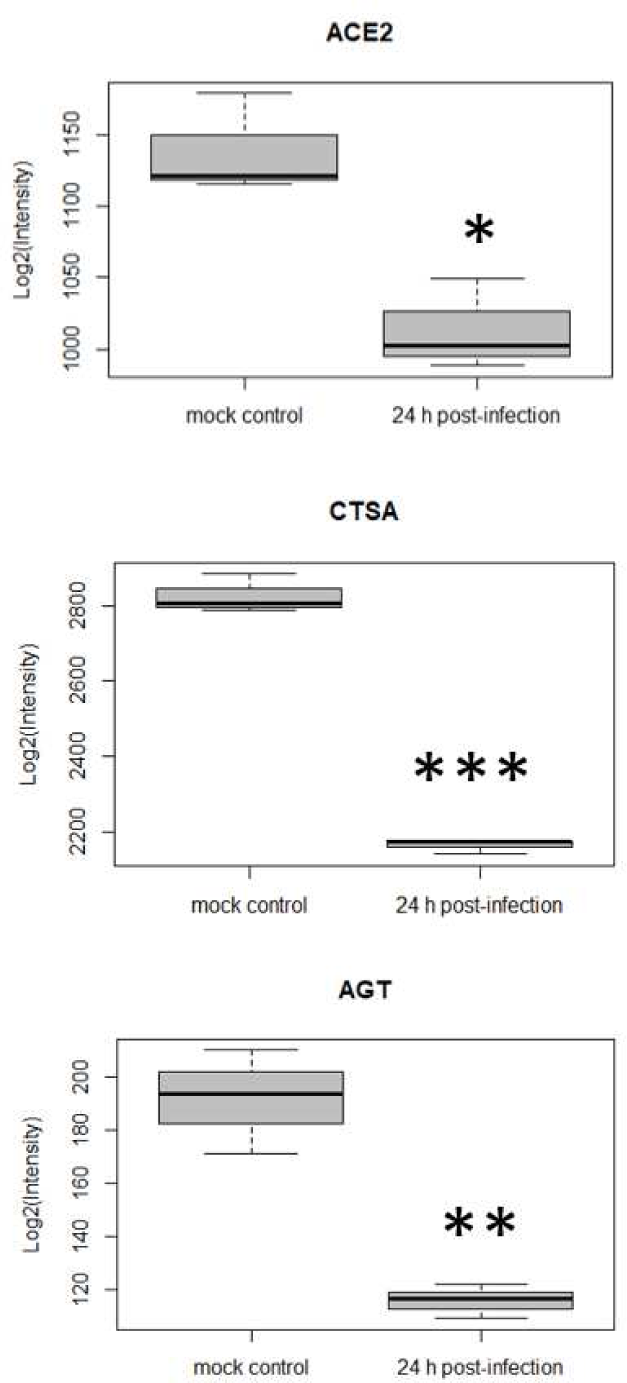
Differential expression for three host cell proteins in the renin-angiotensin system at 24 h post SARS-CoV-2 infection (*p-value<0.05, **p-value<0.01, ***p-value<0.0001, comparison to mock control).

Summarizing, in this work, through a re-analysis of data from a study of the changes caused by SARS-CoV-2 infection in a cellular model, we point out several proteins, mainly related to inflammatory response, but also another subset related to chromosome segregation, that might be being modulated by the infection. In the case of the proteins related to inflammation, the up-regulation observed could be linked to the propagation of inflammatory reaction and lung injury that is observed in advanced stages of COVID-19 patients.

